# Aerobic removal of microcystin-LR by a novel native effective bacterial community designated as YFMCD4 isolated from Lake Taihu

**DOI:** 10.1101/347088

**Authors:** Fei Yang, Jian Guo, Feiyu Huang, Isaac Yaw Massey, Ruixue Huang, Ping Ding, Weiming Zeng

## Abstract

Microcystins (MCs) are a group of monocyclic heptapeptide hepatotoxins produced by species of cyanobacteria. MC-LR is the most toxic and frequently detected MCs variant in water, which poses a great threat to the natural ecosystem and public health. It’s important to seek environment-friendly and cost-efficient methods to remove MC-LR. To investigate the MC-degrading capacities of a novel indigenous bacterial community designated as YFMCD4 and the influence of environmental factors including various temperatures, MC concentrations and pH on the MC-degrading activities, the concentration of MC-LR was measured by high performance liquid chromatography. In addition, the MC-degrading mechanism containing the degradation pathway and products of YFMCD4 was studied using HPLC coupled with an ultra-high resolution LTQ Orbitrap Velos Pro ETD mass spectrometry equipped with electrospray ionization interface. The data showed MC-LR can be removed at the maximum rate of 0.5 µg/(ml·h) by YFMCD4 containing *Alcaligenes faecalis* and *Stenotrophomonas acidaminiohila*. The MC-degrading rates of YFMCD4 were significantly affected by different temperatures, pH and MC-LR concentrations. Two intermediates of a tetrapeptide and Adda appeared in the degradation process. These results illustrate that the novel bacterial community YFMCD4 can remove MC-LR effectively and completely, which indicates YFMCD4 possesses a significant potential to be used in bioremediation of water bodies contaminated by MC-LR.

## Introduction

Cyanobacterial harmful algal blooms (CyanoHABs) have proliferated worldwide because of eutrophication and climate change [1-4]. Microcystins (MCs) produced by *Microcystis*, *Anabaena*, *Oscillatoria* and *Nostoc* during CyanoHABs theats the public health and have become a serious global problem due to their extreme toxicities, which have attracted global attention [3, 5]. MCs are a group of monocyclic heptapeptide hepatotoxins with the common genetic structure cyclo-(D-Ala-X-D-MeAsp-Z-Adda-D-Glu-Mdha-), where X and Z represent variable L-amino acids, and Adda is the b-amino acid residue of 3-amino-9-methoxy-2,6,8-trimethyl-10-phenyldeca-4,6-dienoic acid. Until now, over 100 analogs of MCs have been identified and MC-LR is the most toxic and abundant MC variant [6, 7]. MC-LR is harmful to different organs including liver, intestine, colon, brain, kidney, lung, heart and reproductive system because it can inhibit the activities of protein phosphatases and affect the regulation of miRNA expression in these systems [8, 9]. Even the chronic exposure to low concentrations of MCs also can promote tumor growth. The International Agency for Research on Cancer (IARC) has classified MC-LR as a possible carcinogen because of its potential carcinogenic activity [10]. To reduce MC-LR risks, the World Health Organization (WHO) has proposed a provisional guideline of 1 µg/L MCs in drinking water and this guideline level has been adopted in legislation in many countries such as South America, Australasia, Europe and China [11].

MC-LR is very stable and resistant to many natural factors including extreme pH, high temperature and sunlight in the environment owing to the cyclic structure [3, 6, 12]. Moreover, MC-LR can be accumulated in aquatic organisms and food crops representing a health hazard to human and animals through food chains [13]. It is very important to reduce MC-LR concentration in freshwater ecosystem. However, conventional drinking water treatments have limited efficacy in removing MC-LR. Some physical and chemical methods containing ozonation, chlorination, photocatalysis and electrolysis have been proposed for MC-LR elimination from drinking water. However, all these methods have certain limitations in terms of high operating costs, low efficacy and harmful by-products [3, 6, 14, 15]. It’s desirable that investigators seek other environmentally-benign and cost-efficient methods and technologies to remove MC-LR found in water bodies [3, 6, 14-17].

Several previous investigations demonstrated that microbial biodegradation may be one of the most environmentally-friendly, effective and promising treatment methods for removing MC-LR in natural waters, since it can detoxify MC-LR and don’t generated any apparent potential harmful by-products [3, 6, 14-17]. A few MC-degrading pure bacterial strains have been isolated, identified and had their mechanisms reported, and most of the isolated MC-degrading bacteria were limited to the family *Sphingomonadaceae* [16-18]. In practice, native bacterial communities (indigenous bacterial mixed culture) may be more suitable for degrading MC-LR in the environment compared to the single pure bacterial strains [6,19]. Therefore, it’s very interesting and important to obtain some native mixed bacterial communities for MC-LR removal.

Lake Taihu is the third largest lake with a total water surface area of about 2,338 km^2^ in China. Lake Taihu is essential to millions of people for drinking water, aquaculture, industrial activities, and recreation, but it has experienced CyanoHABs every year during the last three decades [3, 6, 15, 18]. The MCs and odorous during CyanoHABs resulted in more than 2 million residents in Wuxi City being without drinking water for a week. Thus, it’s desirable to obtain bacterial stains and remove MC-LR in water. In this study, the MC-LR removal capacities of a novel native bacterial community designated as YFMCD4 from Lake Taihu were determined under various environmental factors containing different temperatures and pH as well as MC-LR concentrations. Moreover, the MC-removal mechanism including degradation pathway and products of YFMCD4 was investigated.

## Materials and Methods

### Materials and Reagents

MC-LR was purchased from Alexis Corporation and stored −20°C (purity≥95 %). Formic acid and methanol used for high performance liquid chromatography (HPLC) and ultra-high resolution LTQ Orbitrap Velos Pro ETD mass spectrometry equipped with electrospray ionization interface (HPLC-ESI-MS) analysis were purchased from Dikma Technology Incorporation in USA. The mineral salt medium (MSM) for bacterial culture, acquisition and MC-LR removal was prepared as previous study [3, 6, 15]

Acquisition of a novel native bacterial community YFMCD4, isolation and identification of bacterial strains in the bacterial mixed culture

5 g of wet sludge sample was collected from Lake Taihu and suspended in 45 ml MSM. A novel MC-degrading bacterial community was obtained and designated as YFMCD4 in 24 days under the conditions previously described by Yang et al. [6]. The bacterial community YFMCD4 serially diluted with sterile MSM and 0.1 ml of each dilution were inoculated onto nutrient agar (2% agar) plates. Two pure bacterial strains named YFMCD4-1 and YFMCD4-2 were isolated.

16S rRNA gene fragments of YFMCD4-1 and YFMCD4-2 were amplified using PCR with the universal primers 5’-AGAGTTTGATCMTGGCTCAG-3’ and 5’-TACGGYTACCTTGTTACGAACTT-3’) under the conditions previously described by [4]. The PCR products were sequenced by the Sangon Biotech Incorporation located in Shanghai, China. Nucleotide sequences comparisons were conducted using the National Center for Biotechnology Information (NCBI) database (http://www.ncbi.nlm.nih.gov/BLAST).The program ClustalW 2.1 was applied to align the entire similar 16S rRNA gene sequences downloaded from the NCBI database. Phylogenetic trees were successfully generated via the neighbor-joining method using the MEGA software Tamura et al. [20].

### MC-LR degradation by bacterial community YFMCD4

To study MC-LR-degrading ability of YMCD4, the bacterial community YFMCD4 was cultured with MC-LR under different incubation conditions including different temperatures at 20°C, 30°C or 40°C, at MC-LR concentrations 1, 2, 3, 4or 5 µg/ml, and at pH 3, 5, 7, 9 or 11. 100µl samples were withdrawn at intervals and centrifuged (12,000×g, 15 min, 4°C) for monitoring the concentrations of MC-LR in all the samples using HPLC. All the experiments were duplicated with bacterial free samples serving as the control.

### Analysis of MC-LR and its degrading products

The Agilent 1100 HPLC machine with a Zorbax Extend C_18_ column (4.6 × 150 mm, 5 µm, Agilent, USA) and a variable wavelength detector (VWD) set at 238 nm was employed for analyzing MC-LR and degradation products. The mobile phase was a mixture of 0.1 % trifluoroacetic acid aqueous solution and methanol (37:63, v/v) set at a flow rate of 0.8 ml/min, injection volume 10µl and column temperature 40°C.

The MC-degrading products were identified by HPLCcoupled with an ultra-high resolution LTQ Orbitrap Velos Pro ETD mass spectrometry (Thermo Scientific, Germany) equipped with electrospray ionization interface (HPLC-ESI-MS). Both the auxiliary and sheath gases were nitrogen at a flow rate of 30 and 5 psi, respectively. The dry gas temperature was set at 350 °C, and nebulizer pressure at 45 psi. Spectra were recorded in positive modes at a spray voltage of 3.5 kV.

## Results

### Acquisition of bacterial community and identification of bacterial strains

A novel MC-degrading bacterial community named YFMCD4 was obtained. Two bacterial strains designated YFMCD4-1 and YFMCD4-2 were isolated from the bacterial community YFMCD4 and identified according to 16S rRNA gene sequences. YFMCD4-1 and YFMCD4-2 was classified as *Alcaligenes faecalis*and and *Stenotrophomonas acidaminiohila* respectively (Figure 1). The nucleotide sequences of 16S rRNA genes from YFMCD4-1and YFMCD4-2 were deposited in the NCBI database with accession number MH106702 and MH106704 respectively

**Figure 1.**
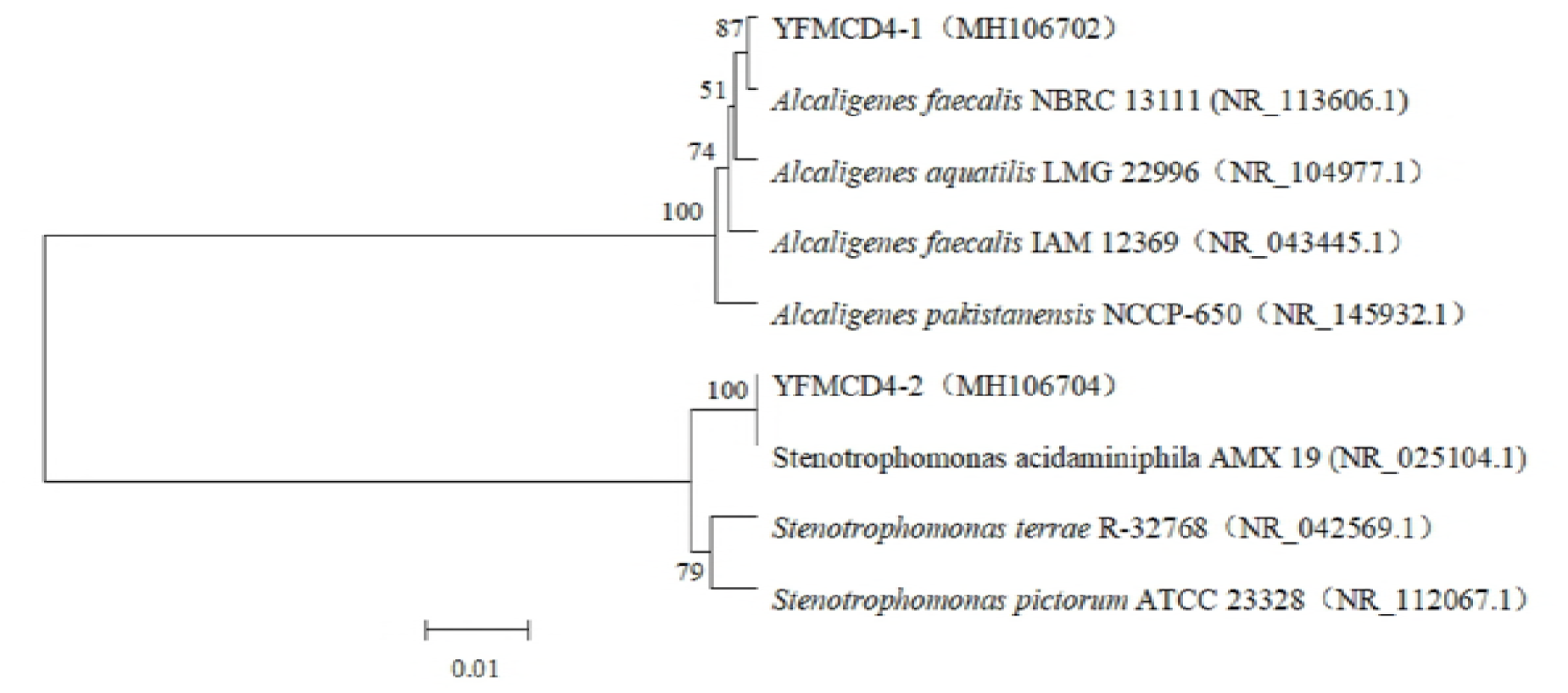
Construction of phylogenetic tree based on the bacterial 16S rRNA gene sequence of the YFMCD4-1 and YFMCD4-2 using neighbor joining method.

### MC-degrading activities under different conditions

Single factor experiments were performed and results are showed in Figures 2-4. The MC-LR degrading rates of the bacterial community YFMCD4 were influenced by different incubation temperatures (Figure 2), MC-LR concentrations (Figure 3) and pH (Figure 4). Figure 2 showed that YFMCD4 degraded MC-LR at the average rate of 0.09, 0.33, 0.25µg/(ml·h) at 20, 30, and 40°C respectively in 10 h.

**Figure 2.**
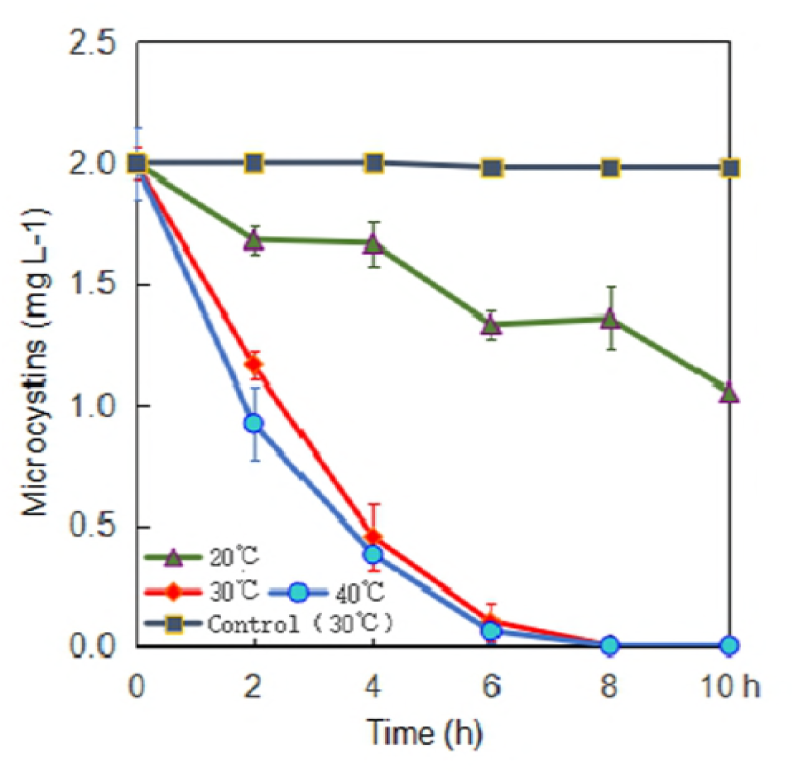
Effect of incubation temperature on the degradation rate of MC-LR by YFMCD4.

**Figure 3.**
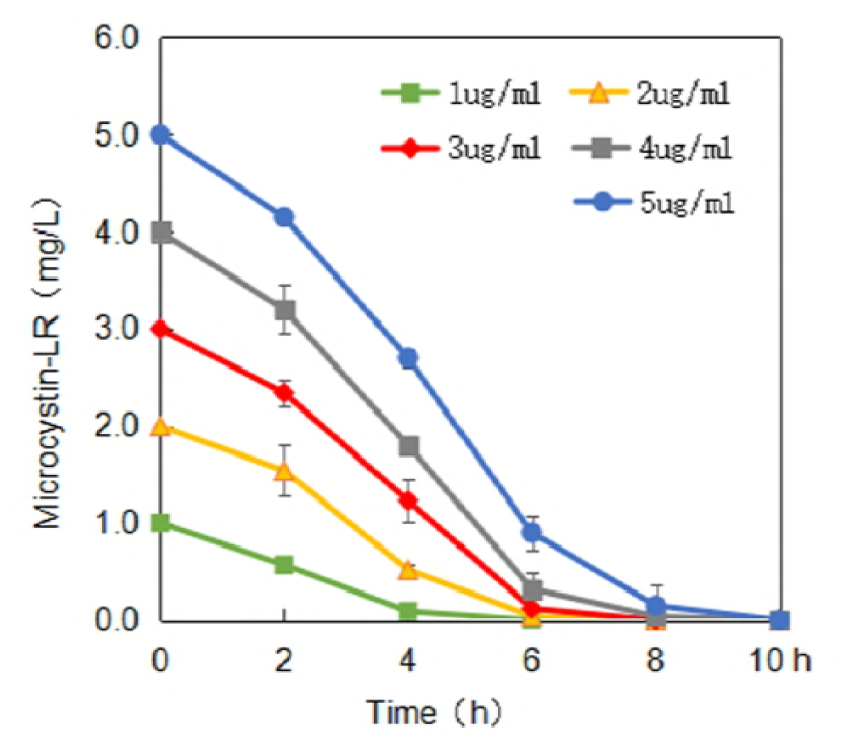
Effect of MC-LR concentration on the degradation of MC-LR by YFMCD4.

**Figure 4.**
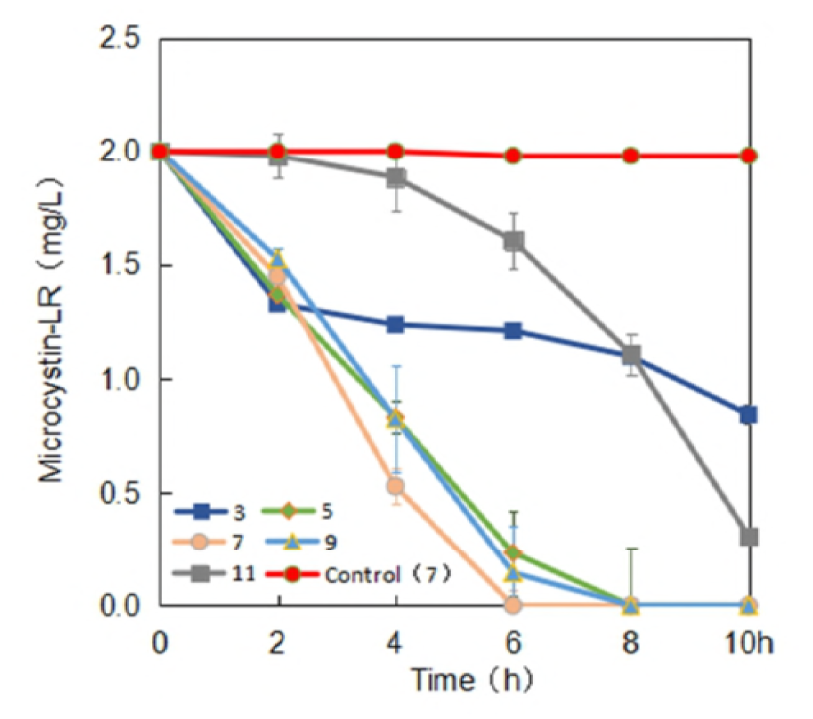
Effect of pH on the degradation of MC-LR by bacterial community YFMCD4.

Figure 3 illustrated that pH 7 and 30°C MC-LR at concentrations of 1, 2, 3, 4 or 5 µg/ml were degraded at the average rate of 0.25, 0.33, 0.375, 0.5 and 0.5 µg/(ml·h) in 10 h, respectively. Figure 4 demonstrated that at 30°C 2 µg/ml MC-LR was degraded by YFMCD4 at the rate of 0. 12, 0.25, 0.33, 0.25 and 0.17 µg/(ml·h) at pH 3, 5, 7, 9 and 11 in 10 h, respectively. Results indicated that the highest MC-degrading rate for YFMCD4 was 0.5 µg/(ml·h) at 30°C and pH 7 with MC-LR concentrations of 4 or 5 µg/ml. It should be noted that there was no MC-LR degradation in the control media without bacterial community YFMCD4.

### MC-LR analysis and degradation products

HPLC chromatograms of MC-LR and its degradation products are showed in Figure 5. Degradation of MC-LR by bacterial community YFMCD4 was tested in culture under the optimal conditions of 30°C, pH 7, with 5 µg/ml of MC-LR concentration in the culture. HPLC chromatograms showed the retention time of MC-LR was 8.1 min (Figure 5a). The peak area of MC-LR decreased significantly after incubation, and two main intermediate degradation products of MC (peak A and B) were apparent at 4h (Figure 5b). The disappearance of all the peaks demonstrated complete catabolism of MC-LR and its degradation products by YFMCD4 in 12 h (Figure 5c). The degradation products peak A and B were further identified using the HPLC-ESI-MS, and exhibited accompanying ion at m/z 615.33850 (Figure 6) and m/z 332.33325 (Figure 7). The HPLC chromatograms and ion of peak A were identical to the tetrapeptide found by [21]. The ion of peak B was identical to Adda which was the final MC degradation product of the *Sphingopyxis* C-1 [22] and the immediate degradation products of *Bordetella* sp. MC-LTH1 [3]. The degradation products indicate that the degradation pathway of YFMCD4 probably may be similar with that of *Bordetella* sp. MC-LTH1 (Figure 8).

**Figure 5.**
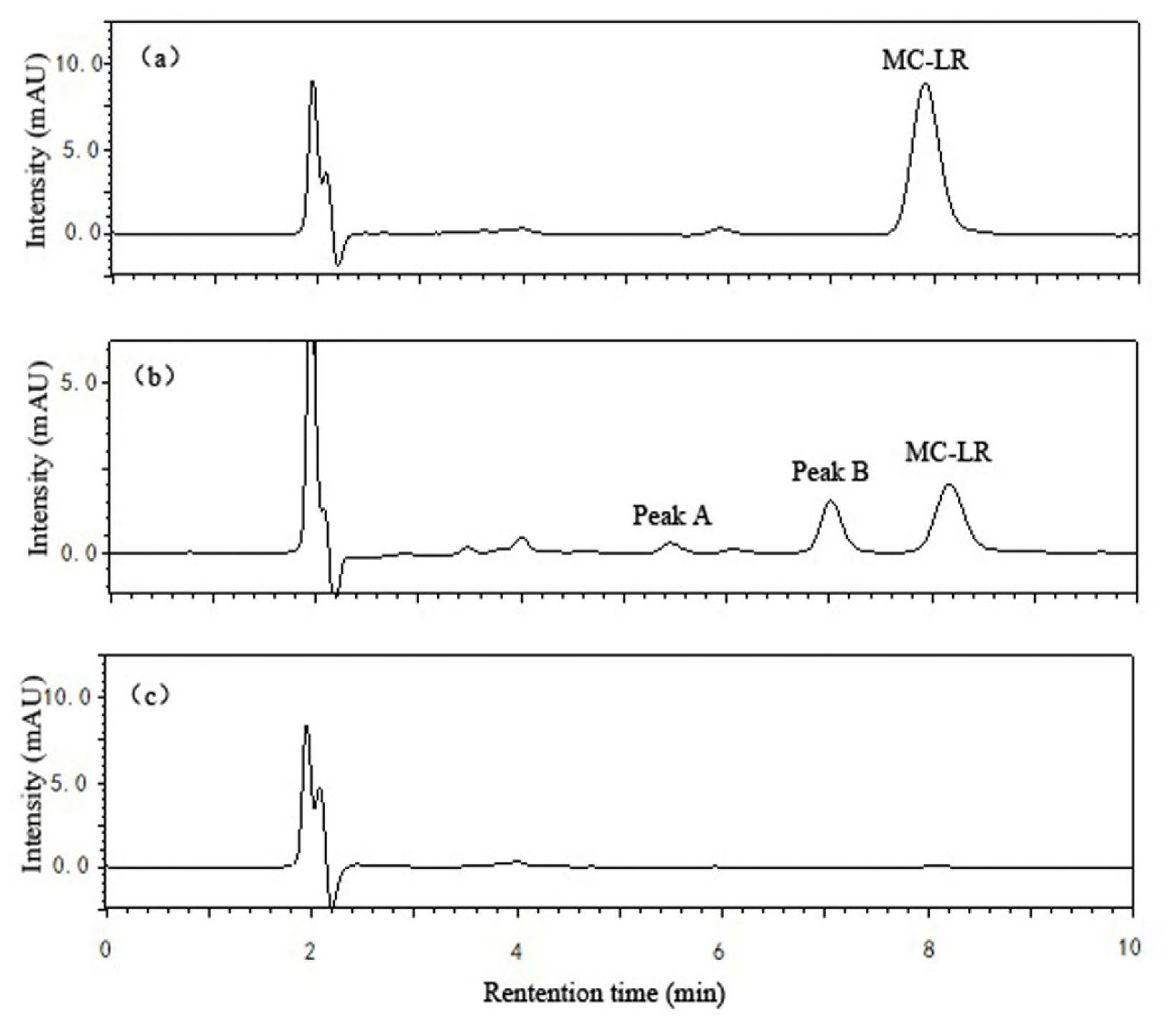
HPLC chromatograms obtained during MC-LR degradation incubated with bacterial community YFMCD4 at time 0 h (a), 4 h (b) and 12 h (c).

**Figure 6.**
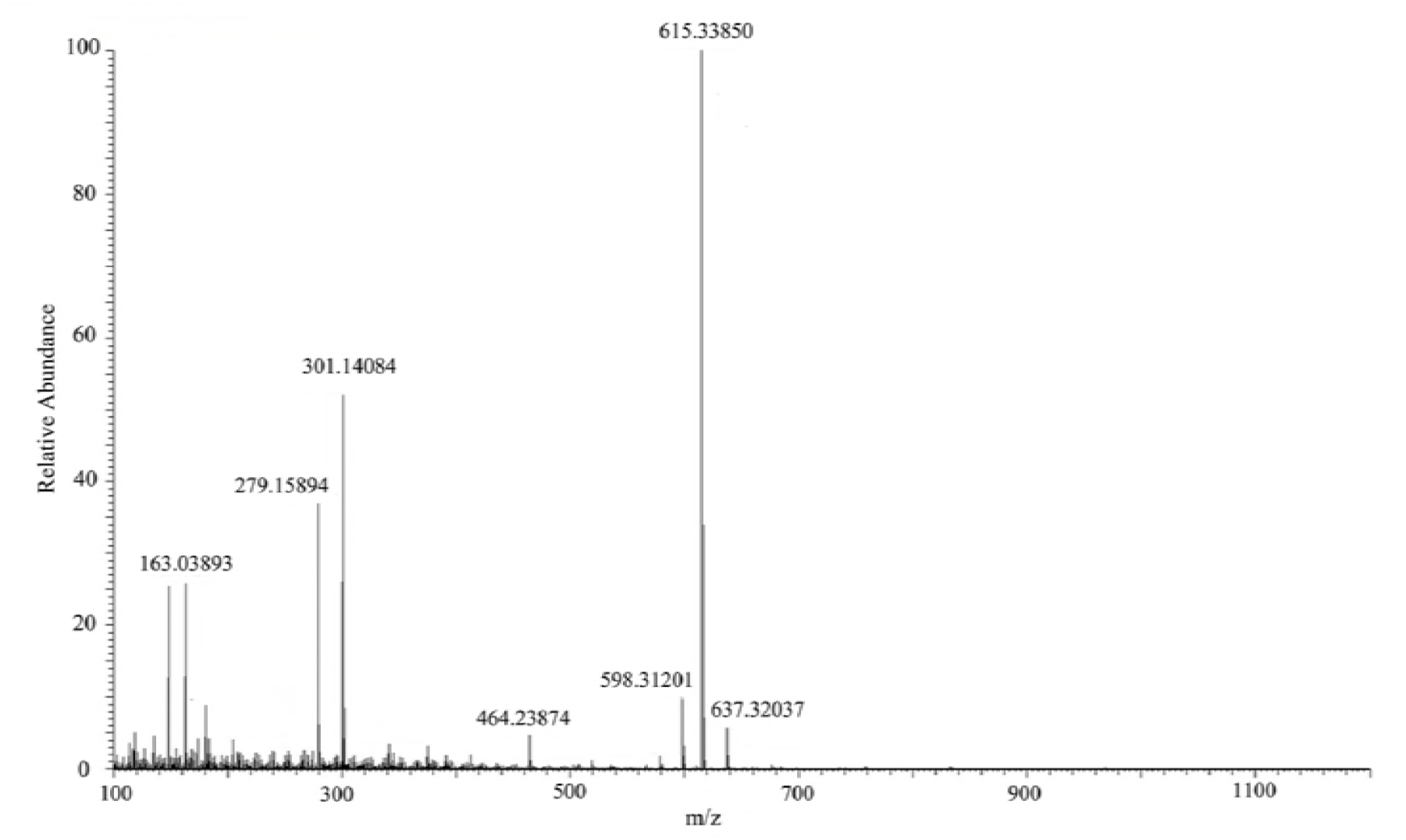
HPLC-E5I-MS spectrum of the biodegradation product A of MC-LR.

**Figure 7.**
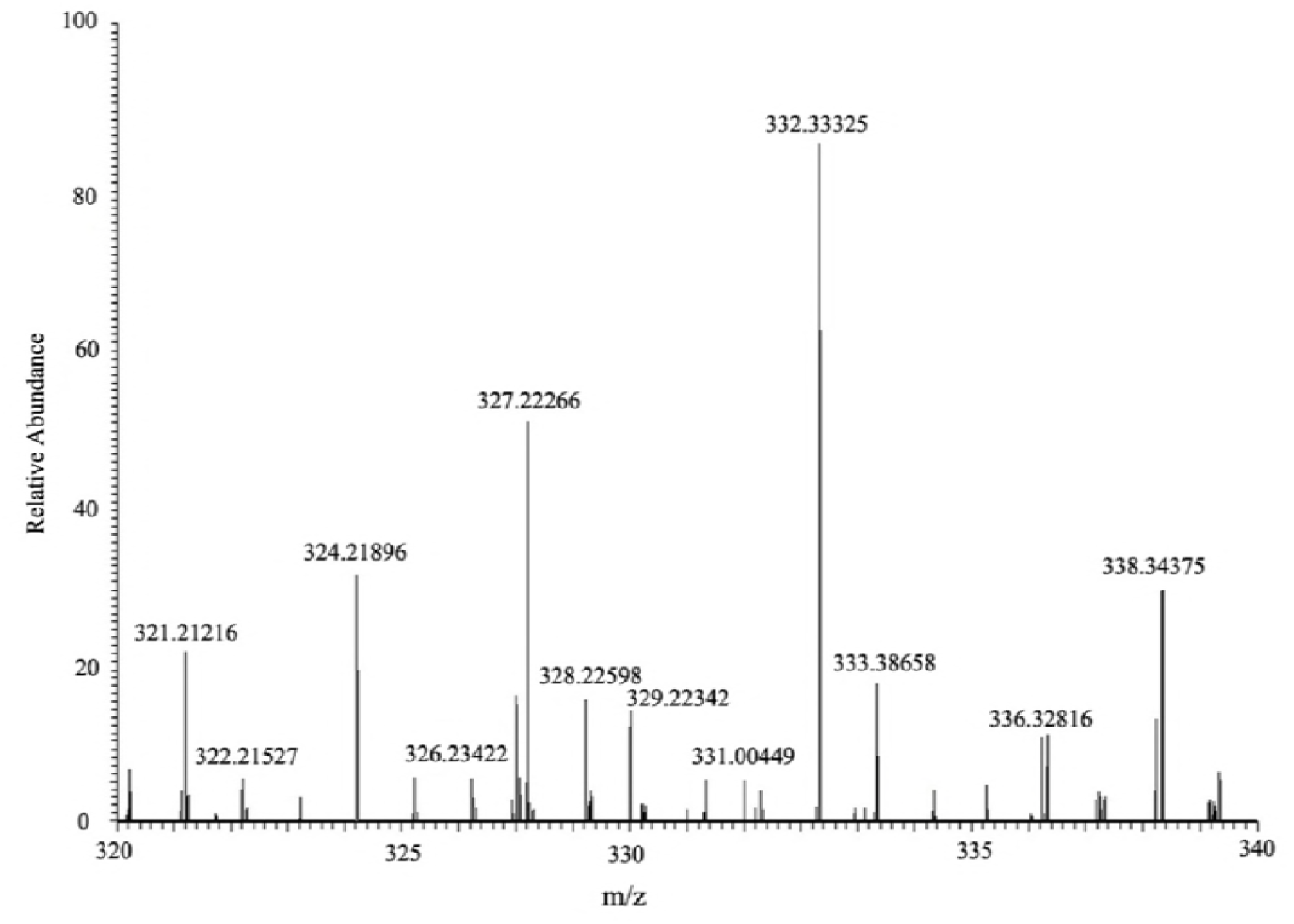
HPLC-ESI-MS spectrum of the biodegradation product B of MC-LR.

**Figure 8.**
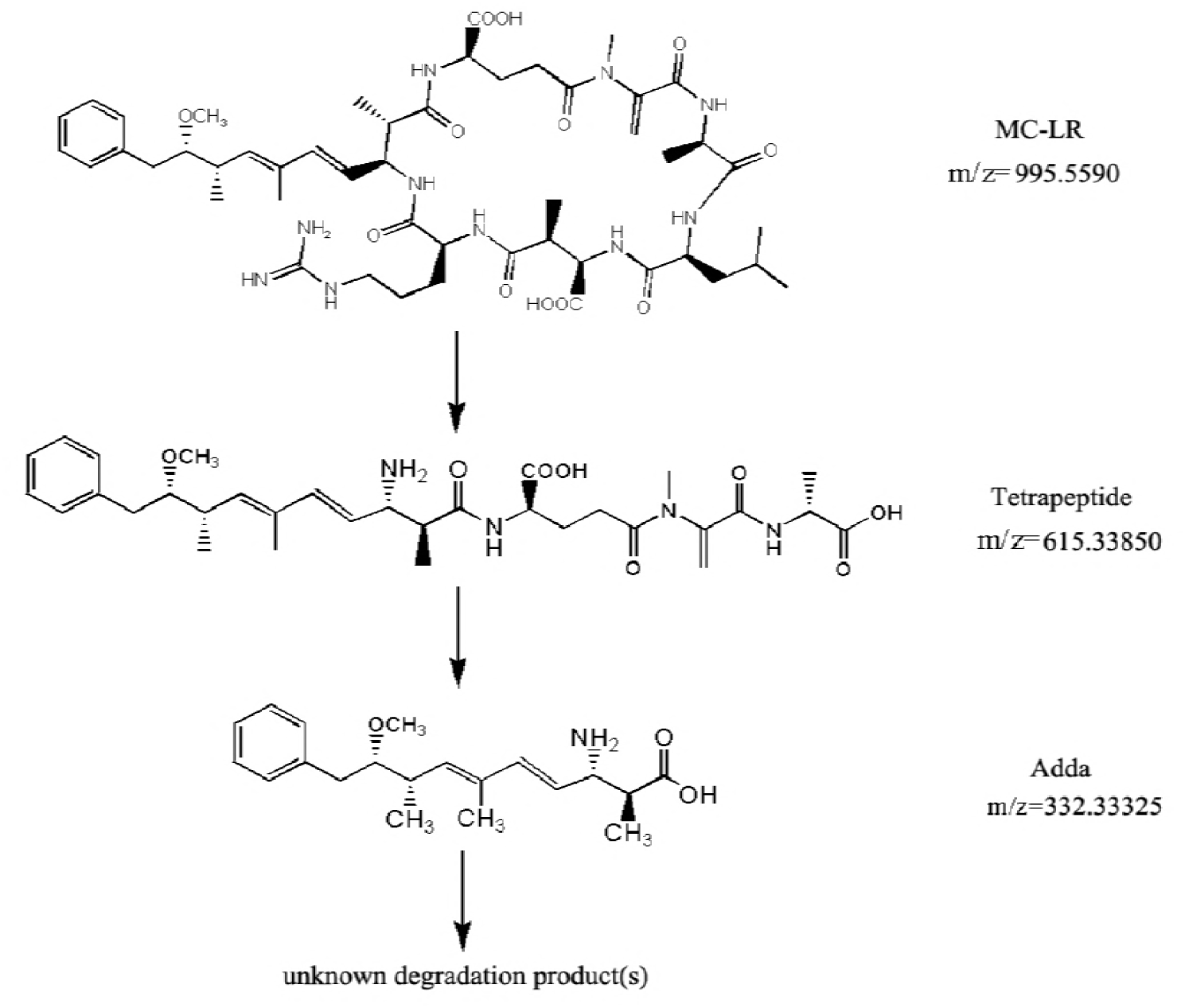
Putative degradation pathway of MC-LR and the formation of intermediate products (tetrapeptide and Adda) by YFMCD4.

## Discussion

Microbial biodegradation is an environmentally-friendly and effective treatment method for detoxify MC-LR in natural waters without potential harmful by-products. Some MC-degrading pure bacterial strains have been isolated and have their MC-LR-degrading rates reported [6, 16, 17, 19]. For example, the single pure bacterial strain *Sphingomonas* sp. ACM-3962 (1.7µg/(ml·d)) [23], LH21 (2.1µg/(ml·d)) [24] and EMS (0.7µg/(ml·d)) [25], *Ralstonia solanacearum* (9.4µg/(ml·d)) [26], *Bordetella* sp. MC-LTH1 (7.4µg/(ml·d)) [3] and *Stenotrophomonas* sp.MC-LTH2 (3 µg/(ml·d)) [15] were studied. Until now, only a few MC-degrading bacteria mixed cultures have been obtained and investigated [16, 27]. Cousins et al. [27] reported that a bacterial community showed the MC-degrading rate of 1.4 × 10^−3^ µg/(ml·d) while what kinds of bacteria existed in the community still needed to be further studied. Ramani et al. [16] found a bacterial community containing two pure bacterial strains *Rhizobium* sp. DC7 and *Microbacterium* sp.DC8 degraded MC at 0.18 µg/(ml·d) while individual DC7 or DC8 can’t degrade MC-LR respectively. Tsao et al. [19] discovered a mixed culture with MC-degradation rate of 0.876 µg/(ml·d), which contained *Sphingomonas* sp., *Pseudoxanthomonas* sp., *Hyphomicrobium aestuarii*, *Sphingobium* sp., *Rhizobium* sp., *Steroidobacter* sp. and *Acinetobacter* sp. Yang et al. [6] declared another natural bacterial community YFMCD1 including *Klebsiella* sp. YFMCD1-1 or *Stenotrophomonas* sp. YFMCD1-2 with the MC-degrading rate of 12 µg/(ml · d). The bacterial community YFMCD4 showed a higher degradation rate of MC-LR at 12 µg/(ml·d) compared with the single bacterial strain and most prior bacterial communities. These results confirmed that indigenous bacterial community always appeared to be more effective and suitable for degrading MC-LR than single pure bacterial strains, which was in accordance with the previous findings by Yang et al. [6], Ramani et al. [16] and Zhang et al. [26]. In general, different bacterial community owed different MC-degrading rates because they contained different pure bacterial species. However, it’s interesting that the bacterial community YFMCD4 exhibited the same highest MC-degrading rate with YFMCD1, which means that different bacterial communities may have a similar MC-degrading rate although they consisted of different kinds of pure bacterial strains [3, 15].

The MC-degrading rates of bacterial community YFMCD4 were significantly affected by different temperatures, pH and MC-LR concentrations. The optimal conditions for degradation of MC-LR by YFMCD4 occurred at 30°C, pH 7 and MC-LR concentration of 4 µg/ml or 5 µg/ml. Prior studies and our previous studies showed that these three factors play an important role in MCs degradation [15, 28-30]. Park et al. [29] found the degradation rates were strongly dependent on temperature and the MC-degrading rate was very slow at 5°C while the maximum degradation rate occurred at 30°C. Ramani et al. [16] found temperature has some effects on MC-LR degradation and the optimal degradation rate was achieved at 26°C. Yang et al. [6] also discovered that the degradation rates changed when the temperature varied, and the best temperature for MC-LR degradation was 30°C. It was necessary to investigate the influence of pH on MC-LR degradation activities of bacteria because the pH of water bodies varies during cyanobacterial blooms [31]. The highest ability of YFMCD4 to degrade MC-LR under neutral environment suggested that YFMCD4 may contain MC-degrading enzymes which are different from alkali-tolerant protease secreted by *Sphigopyxis* sp. C-1 [31].

Two kinds of intermediates of MC-LR degradation were identified as the linearized MC-LR and a tetrapeptide in the previous studies [3, 15, 21, 32]. Moreover, the intact Adda was isolated and identified from the final MC-LR degradation products using *Sphingomonas* sp. B-9 [32]. In this study, two intermediates Adda and a tetrapeptide appeared when YFMCD4 degraded MC-LR, and the Adda disappeared finally. Therefore, the results showed that, the MC-degrading mechanism of the bacterial community YFMCD4 is different from that of the previous bacteria *Sphingomonas* sp. B-9 [32] and ACM-3962 [23] as well as *Sphigopyxis* sp. C-1 [31]. The MC-degrading mechanism of the bacterial community YFMCD4 is also possibly different from that of the bacterial community YFMCD1 because of no existence of tetrapeptide in the MC-degrading products using YFMCD1. As is known to us all, the Adda is absolutely essential for the biological activities of MC-LR [32]. In this study, the Adda was completely degraded, which suggested that the bacterial community YFMCD4 had the capacity of detoxifying MC-LR [3, 6, 15]. The degradation products of Adda needed to be further isolated and clarified, and it is important to investigate the practical MC-degrading effects of YFMCD4 when it is applied into different kinds of water polluted by MC-LR in the future.

## Conclusion

A novel native effective MC-degrading bacterial community designated as YFMCD4 was obtained from Lake Taihu, and two pure bacterial strains *Alcaligenes faecalis* YFMCD4-1and *Stenotrophomonas acidaminiohila* YFMCD4-2 were isolated from the bacterial community YFMCD4. The degradation rate of MC-LR by bacterial community YFMCD4 was significantly influenced by various pH, temperature and MC-LR concentrations, and the highest rate reached 0.5 µg/(ml·h) at 30°C and pH 7 with MC-LR concentrations of 4 or 5 µg/ml. Two intermediates of tetrapeptide and Adda existed in the MC-degrading products, and the Adda was completely degraded by the bacterial community YFMCD4. Therefore, the bacterial community YFMCD4 can completely degrade MC-LR effectively and has a great potential for the bioremediation of water polluted by MC-LR.

## Acknowledgments

We would like to thank Professor Shaogang Liu from Central South University for his help on MC degradation products measurements and Associate Professor Jihua Chen from Central South University for his advice on MC degradation.

